# Lifetime changes in CD4 count, viral load suppression and adherence among adolescents living with HIV in urban Peru

**DOI:** 10.1101/580084

**Authors:** Carly A Rodriguez, Lenka Kolevic, Alicia Ramos, Milagros Wong, Maribel Munoz, Kunjal Patel, Molly F Franke

## Abstract

**Introduction:** Viral load suppression and adherence to combined antiretroviral therapy (cART) have been shown to be lower in adolescents than in other age groups; however, this relationship has not been documented longitudinally from childhood to adolescence and has rarely been examined outside of high-resource settings and sub-Saharan Africa. To address this knowledge gap, we quantified longitudinal changes in CD4 cell count, viral load suppression, and cART adherence in adolescents living with HIV in urban, Peru.

**Methods:** We conducted a retrospective chart review among adolescents ages 10-18 years on cART and receiving care at a large, public sector pediatric hospital as of December 2015. We abstracted clinical notes indicating nonadherence and viral load and CD4 counts from childhood to adolescence. We modeled the association between age and each outcome with restricted cubic splines accounting for multiple observations per person, and graphed study outcomes by age.

**Results:** A median of 7.7 years (25^th^ percentile=4.9, 75^th^ percentile=10.2) of follow up were observed for 128 adolescents. Nearly 70% of patients had at least one nonadherence note and the proportion with nonadherence increased log-linearly with age (p<0.0001). The peak proportion with viral load suppression was 84% (95% CI: 79, 88) at age 13, which dropped to 67% (95% CI: 47, 83) by age 18. Mean CD4 count decreased at age 13, dropping from 723 cells/mm^3^ (95% CI: 666, 784) to 429 cells/mm^3^ (95% CI: 356, 517) by age 18.

**Conclusion:** This is the first report from Latin America to examine longitudinal changes in HIV outcomes from childhood into adolescence. Consistent with the limited evidence from other settings, decreases in viral load suppression and mean CD4 count occurred in early adolescence in tandem with increases in nonadherence. Adolescent-friendly cART adherence support interventions to target this critical period are urgently needed.

## Introduction

Combined antiretroviral therapy (cART) has greatly improved survival for infants and children perinatally infected with HIV. As a result, adolescents make up a growing portion of the global HIV burden with an estimated 1.8 million adolescents living with HIV (ALHIV) between ages 10 and 19 years in 2017.[1] Lower viral load suppression rates have been observed in children and adolescents under 15 years than in adults[2] and uptake and adherence to cART is reported to be lower in adolescents than other age groups.[3-6] Furthermore, HIV-related deaths among adolescents have tripled over the last two decades, occurring primarily in those perinatally infected.[7]

Little is known about longitudinal changes in HIV outcomes across the lifespan, particularly during childhood and adolescence–a period of dramatic physical growth and cognitive development. Available studies on the long-term outcomes of perinatally infected adolescents have short follow-up periods and are largely from high resource settings or Sub Saharan Africa, [8-11] with limited data from cohorts in Latin America.[12,13] The objective of this study was to examine changes in absolute CD4 count, viral load suppression, and adherence from childhood to adolescence among patients on cART in Lima, Peru.

## Methods

### Study design

We conducted a retrospective chart review of ALHIV ages 10 to 18 years receiving care at the Instituto Nacional de Salud del Niño (National Institute for Child Health, henceforth INSN) in Lima, Peru. INSN is a national public sector referral hospital for pediatric care and hosts the largest HIV treatment clinic for children and adolescents in the country. In 2004, cART became widely available in Peru due to the expansion of free, universal access to HIV care and treatment.[14] The earliest guidance for treatment of HIV in children and adolescents in 2003 recommended two nucleoside reverse transcriptase inhibitors (NRTIs) (zidovudine and lamivudine) and one protease inhibitor (nelfinavir).[15] This guidance was updated in 2013 to recommend two NRTIs and one non-nucleoside reverse transcriptase inhibitor (nevirapine or efaviranez).[16] Between June 2015 and April 2016, trained study personnel abstracted demographic data and cART treatment history, including longitudinal CD4 measures, viral load counts, and clinical notes describing nonadherence after cART initiation from paper clinical charts for 132 adolescents. All adolescents were alive and on cART at the time of the chart review.

### Study outcomes

Absolute CD4^+^ lymphocyte monitoring was conducted every three months and viral load monitoring was conducted every six months per national guidelines.[15,16] CD4 count and viral load and measurements <6 months after cART initiation were excluded to allow values to stabilize post-treatment initiation. This led to the exclusion of four adolescents from the analysis who were on cART for <6 months at the time of the chart review. CD4 counts were additionally restricted to ages 5 to 18 years because of the tendency for CD4 counts to be higher at young ages due to age-related immune development.[17] Adherence and viral load were restricted to the same age-years for comparability. Due to varying viral load lower limits of detection over the follow-up period, we defined viral load suppression as <400 copies/ml, the least sensitive threshold. Providers routinely assessed nonadherence using clinical judgement during medical encounters; therefore, the absence of clinician-documented nonadherence at a given encounter was presumed to indicate adequate adherence at that time.

### Statistical analysis

We conducted longitudinal analyses of the three study outcomes using generalized estimating equations and graphed the mean CD4 count and predicted probabilities of viral load suppression and nonadherence by age-month.

In generalized estimating equations for CD4 count we used a normal response and log link; for viral load suppression and nonadherence we used a binary response and logit link. We applied an autoregressive correlation matrix to account for multiple observations per patient.[18,19] The autoregressive correlation matrix assumes measurements closest in time are most correlated and that this correlation decreases exponentially as measurements become more distant. To examine the possible non-linear association between age and each outcome nonparametrically, we used restricted cubic splines. We fit multiple models with varying numbers of knots for each outcome and chose the best model by assessing the quasi-likelihood under the Independence Criterion (QIC), where smaller values indicate better model fit.[20] For the outcomes of CD4 count and viral load suppression, modeling age with seven and four knots, respectively, fit the data best. For the outcome of nonadherence, modeling age as linear fit the data best. We tested for a relationship between age and each outcome using the likelihood ratio test. To examine whether the overall association between age and viral suppression was driven by an increasing probability of viral suppression at younger ages, versus a declining probability of suppression during adolescence, we tested for an association between older age (16-18 years) versus younger age (10-12 years) in the subset of measurements taken from 10 to 18 years of age. Analyses were conducted in SAS version 9.4 (Cary, NC). This study was reviewed and approved by research ethics committees at INSN, Lima, Peru and Harvard Medical School, Boston, USA. Research ethics committees deemed that patient consent was not required for the retrospective chart review.

## Results

### Cohort characteristics

Of 132 adolescents, 128 (97.0%) were on cART for ≥6 months at the time of chart review and were included in the analysis. The median age at the time of the chart review was 14.6 years (25^th^ percentile=12.1, 75^th^ percentile=16.6) with a median follow-up time after six months of cART of 7.5 years (25^th^ percentile=4.8, 75^th^ percentile=10.0). The median age of cART initiation was 5.7 years (25^th^ percentile=3.8, 75^th^ percentile=9.4) (Table 1). Because data were restricted to the period that patients were age five or greater and on cART, the earliest CD4 counts, viral load measurements, and nonadherence notes were recorded in 2004 (i.e. the year in which the oldest patients included in the chart review were age five and had access to cART). The latest CD4 counts, viral load measurements, and nonadherence notes were recorded in 2015 (i.e. the year of the chart review).

**Table 1.** Clinical characteristics among adolescents living with HIV in urban Peru, (N=128)

|  | Median (25<sup>th</sup>, 75<sup>th</sup> percentiles)<sup>a</sup> |
| --- | --- |
| Male, N (%) | 68 (53.1) |
| Age at chart review (years) | 14.6 (12.1, 16.6) |
| Age at cART initiation (years) | 5.7 (3.8, 9.4) |
| Follow-up time on ≥6 months of cART (years) | 7.5 (4.8, 10.0) |
| Time on cART (years) | 8.2 (5.4, 10.7) |
| Viral loads measurements |  |
| Number of viral loads measurements per adolescent after ≥6 months of cART | 12 (9, 16.5) |
| Interval between viral load measurements (days) | 182 (175, 203) |
| CD4 count measurements |  |
| Number of CD4 count measurements per adolescent after ≥6 months of cART | 17 (11, 27) |
| Interval between CD4 counts measurements (days) | 91 (91, 126) |
| Nonadherence |  |
| Adolescents with a nonadherence note, N (%) | 86 (67.2) |
| Number of nonadherence notes per adolescent | 1 (0, 4) |

<sup>a</sup> Unless specified otherwise

### CD4 count

A total of 2,449 CD4 counts after ≥6 months of cART were observed in 128 patients. Patients had a median of 17 CD4 counts (25^th^ percentile=11, 75^th^ percentile=27) over the follow up period and the median interval between measurements was 3.0 months (25^th^ percentile=3.0 months, 75^th^ percentile=4.1 months). We observed a statistically significant non-linear relationship between age and CD4 count (p=0.02, Figure 1). The mean CD4 count trended downward during childhood and decreased at a faster rate after approximately 12.8 years of age (154 months) from a predicted mean CD4 count of 723 cells/mm^3^ (95% CI: 666, 784) to 429 cells/mm^3^ (95% CI: 356, 517) at 18 years of age (216 months).

**Figure 1.**
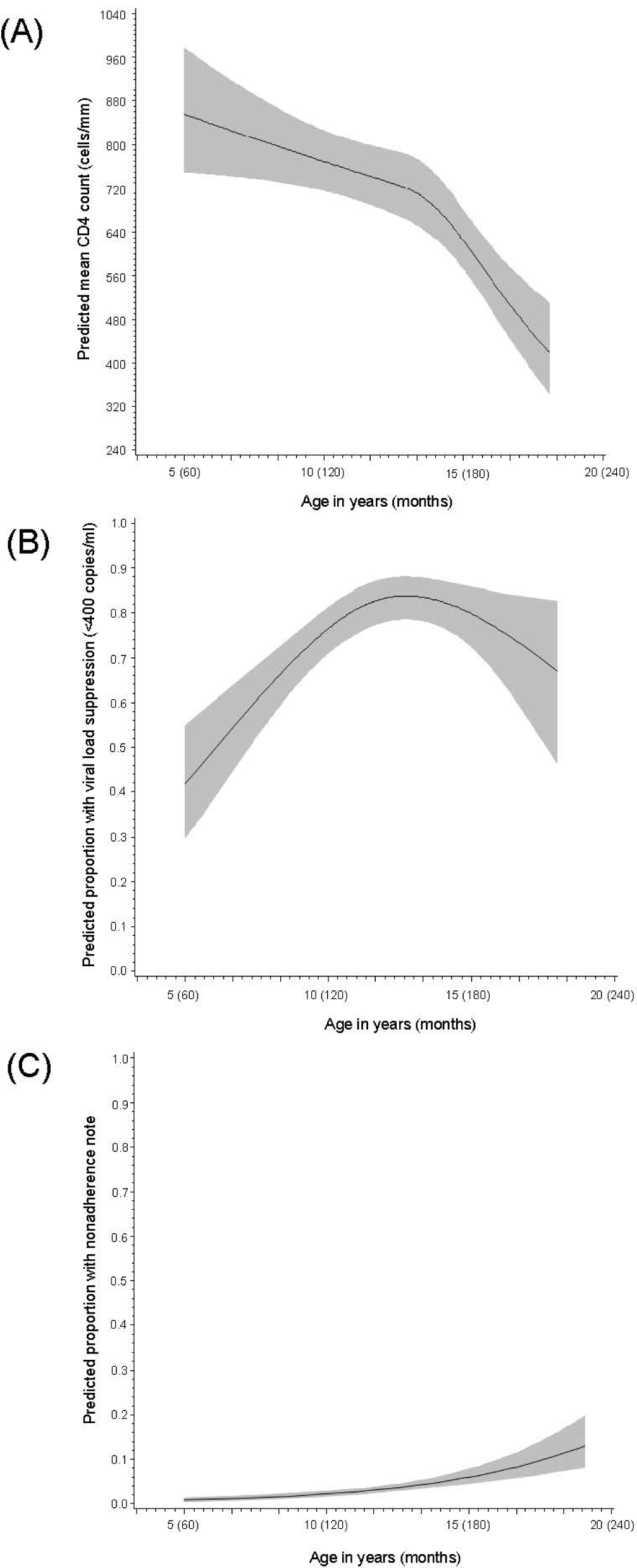
Lifetime changes in CD4 count (A), viral load suppression (B), and nonadherence (C) from age 5 (60 months) to 18 years (216 months) among adolescents living with HIV on cART in Lima, Peru. (A) Mean CD4 count (cell/mm^3^), by age in years and months; (B) Proportion of adolescents with viral load suppression (<400 copies/ml), by age in years and months; (C) Proportion of adolescents with a nonadherence note, by age in years and months

### Viral load suppression

Over the follow up period, patients had a median of 12 viral load measurements after having been on cART for ≥6 months (25^th^ percentile=9, 75^th^ percentile=16.5). The median interval between assessments was 6.0 months (25^th^ percentile=5.8 months, 75^th^ percentile=6.7 months). Of 1,531 viral loads among 128 patients, 1,115 (73%) were suppressed (<400 copies/ml). At least one suppressed viral load was observed in 123 (96%) patients. We observed a non-linear relationship between age and viral load suppression (p<0.0001) in which the predicted proportion of patients with viral suppression increased during childhood and early adolescence and declined thereafter (Figure 1). The predicted proportion of patients with viral load suppression peaked at age 12.7 years (152 months), with 84% (95% CI: 79, 88) virally suppressed. By 18 years of age (216 months), suppression rates decreased to 67% (95% CI: 47, 83). When we examined the association between age and viral load suppression during adolescence, we found a significantly lower predicted probability (p=0.02) of viral load suppression in older adolescence (i.e., 16 to 18 years of age) than younger adolescence (i.e., 10 to 12 years of age).

### Nonadherence

A total of 328 nonadherence notes were recorded over 10,788 follow-up months. Eighty-six (67%) patients had at least one nonadherence note recorded in their chart. The median number of nonadherence notes per patient was one (25^th^ percentile=0, 75^th^ percentile=4). The predicted proportion with a nonadherence note increased with age in a log-linear fashion (p<0.0001, Figure 1). At five years of age (60 months), the predicted proportion of patients with a nonadherence note was 0.8% (95% CI: 0.5, 1.3) which increased to 11% (95% CI: 7, 15) by 18 years of age (216 months).

## Discussion

We report lifetime changes of CD4 count, viral load suppression and nonadherence from childhood into adolescence in a cohort of adolescent patients on cART at an urban pediatric hospital in Lima, Peru. We found that dramatic declines in CD4 count and viral load suppression were observed after approximately age 13 and that nonadherence increased with age. These findings indicate that targeted interventions to improve clinical outcomes and support cART adherence are needed early in adolescence in this population.

Adherence to cART is critical to maintaining viral load suppression.[21,22] In our study, decreases in viral load suppression and mean absolute CD4 count with age are likely due to increases in nonadherence, which changed in tandem with these outcomes. Other work in this population supports that adherence suffers during adolescence and that the mechanisms through which nonadherence occur are amenable to intervention. Through a health behaviors survey implemented in this study population, we found self-reported nonadherence was greatest in the 13 to 15 year age group, with 82% of adolescents missing ≥3 doses in a 30 day period.[23] In psychosocial support groups, adolescents described barriers to adherence at the individual-and family/caregiver-levels, providing the ideal opportunity to deliver support or education interventions.[24] Over the last decade, many adherence interventions leveraging technology have been studied.[25,26] Some of this work has been conducted exclusively in adolescents[27,28] and may be particularly acceptable in populations with ready access and experience using mobile devices, like adolescents in Lima.[29] As adolescents mature and responsibility for their care is transferred from caregiver to the adolescent, health education on living with HIV becomes increasingly important. Education through non-traditional mediums such as social media and music have been explored in young persons,[30-32] however further work from a variety of settings is needed. At 18, most adolescents must transition from pediatric to adult HIV care, a period associated with poor outcomes.[33] In our study, we observed declines in viral load suppression and adherence even before this transition. In anticipation of this change, education and skills building on how to stay healthy into adulthood should be provided early on in adolescence.

Adolescents continue to be underrepresented in HIV research and policy, despite calls to prioritize this group.[34,35] Systematic reviews assessing interventions for ALHIV have found that most studies are conducted in adults or in high-resource settings.[36-38] When studies include ALHIV from low-resource settings, they are primarily from Sub Saharan Africa,[38] limiting the generalizability of findings outside settings with generalized HIV epidemics.[38] A 2015 systematic review of interventions to improve linkage and retention in care among ALHIV did not identify any studies from Latin America,[27] and only 10% of studies in a review investigating adherence among adolescents were conducted in Latin America—all of which were from Brazil.[38] In Peru, adolescents ages 10 to 19 years are largely perinatally infected, while new cases of HIV in youth ages 15 to 24 years are concentrated in men who have sex with men and transgender women.[39] These key differences in risk populations demand tailored interventions.

Limitations include that our study population is a survivor cohort. A retrospective chart review of all children <18 years receiving HIV care at INSN from 2003 to 2012 reported mortality rates were under 9%, indicating survivor bias in our study is likely small.[40] Additionally, data on nonadherence was based on the presence of a clinician note documented in the chart. In modeling these data, we assumed that the absence of a nonadherence note signified adherence, thus our estimate of the predicted proportion of patients with a nonadherence note may be of greater magnitude. Despite these limitations, the nonadherence results triangulate with those of CD4 count and viral load, as would be expected. Together, these findings provide evidence that adolescence is an important period for interventions aimed at improving clinical outcomes, of which adherence support is one strategy.

## Conclusions

Studies assessing lifetime HIV outcomes from childhood to adolescence are limited, especially in Latin America. Decreases in mean CD4 count and viral load suppression were observed during early adolescence and occurred in accordance with provider-reported nonadherence. Research on effective, tailored interventions aimed at improving clinical outcomes and adherence during adolescence are needed in this population.

## Authors’ contributions

LK, MFF and MM conceived of the study. MFF designed the study. AR and MW collected the data and MM supervised data collection. CAR conducted the analysis and wrote the first draft. AR, CAR, KP, LK, MFF and MW interpreted the results. All authors critically reviewed the manuscript and approved the manuscript for submission.

## Acknowledgements

This study was support by funding from the William F. Milton Fund of Harvard University.

## References

1. UNICEF. Turning the tide against AIDS will require more concentrated focus on adolescents and young people. 2018. Accessible at: https://data.unicef.org/topic/hivaids/adolescents-young-people/. Accessed September 17, 2018.

2. Boerma RS, Boender TS, Bussink AP, Calis JCJ, Bertagnolio S, Rinke de Wit TF, et al. Suboptimal Viral Suppression Rates Among HIV-Infected Children in Low-and Middle-Income Countries: A Meta-analysis. Clin Infect Dis. 2016;63(12):1645–54.

3. Wringe A, Floyd S, Kazooba P, Mushati P, Baisley K, Urassa M, et al. Antiretroviral therapy uptake and coverage in four HIV community cohort studies in sub-Saharan Africa. Trop Med Int Health. 2012;17(8):e38–48.

4. Children and AIDS Fifth stocktaking report. New York, United Nations Children’s Fund, 2010. Accessible at: http://www.unicef.org/publications/files/Children_and_AIDSFifth_Stocktaking_Report_2010_EN.pdf. Accessed September 17, 2018.

5. Williams PL, Storm D, Montepiedra G, Nichols S, Kammerer B, Sirois PA, et al. Predictors of adherence to antiretroviral medications in children and adolescents with HIV infection. Pediatrics 2006; 118:e1745–57.

6. Khan M, Song X, Williams K, Bright K, Sill A, Rakhmanina N. Evaluating adherence to medication in children and adolescents with HIV. Arch Dis Child 2009; 94:970 LP–973.

7. UNAIDS/UNICEF. All in to end the adolescents AIDS epidemic. 2016. Accessible at: http://www.unaids.org/sites/default/files/media_asset/ALLIN2016ProgressReport_en.pdf. Accessed September 17, 2018.

8. Dollfus C, Le Chenadec J, Faye A, Blanche S, Briand N, Rouzioux C, et al. Long-term outcomes in adolescents perinatally infected with HIV-1 and followed up since birth in the French perinatal cohort (EPF/ANRS CO10). Clin Infect Dis. 2010;51(2):214–24.

9. Neilan AM, Karalius B, Patel K, Van Dyke RB, Abzug MJ, Agwu AL, et al. Association of Risk of Viremia, Immunosuppression, Serious Clinical Events, and Mortality With Increasing Age in Perinatally Human Immunodeficiency Virus-Infected Youth. JAMA Pediatr. 2017;171(5):450–60.

10. Smith TT, Hsu AJ, Hutton N, Womble F, Agwu AL. Long-term Virologic Suppression Despite Presence of Resistance-associated Mutations Among Perinatally HIV-infected Youth. Pediatr Infect Dis J. 2015;34(12):1365–8.

11. Patel K, Hernan MA, Williams PL, Seeger JD, McIntosh K, Van Dyke RB, et al. Long-term effectiveness of highly active antiretroviral therapy on the survival of children and adolescents with HIV infection: a 10-year follow-up study. Clin Infect Dis. 2008;46(4):507–15.

12. Souza E, Santos N, Valentini S, Silva G, Falbo A. Long-term follow-up outcomes of perinatally HIV-infected adolescents: infection control but school failure. J Trop Pediatr. 2010;56(6):421–6.

13. Candiani TMS, Pinto J, Cardoso CAA, Carvalho IR, Dias ACM, Carneiro M, et al. Impact of highly active antiretroviral therapy (HAART) on the incidence of opportunistic infections, hospitalizations and mortality among children and adolescents living with HIV/AIDS in Belo Horizonte, Minas Gerais State, Brazil. Cad Saude Publica. 2007;23 Suppl 3:S414–23.

14. Ministerio de Salud. A Step Forward in the Fight Against AIDS In Lima: Ministerio de Salud del Peru (MINSA): The first two years of universal access to antiretroviral treatment in Peru. Lima, Peru, 2006.

15. Ministerio de Salud. Resolución Ministerial N° 731-2003-SA/DM, que aprueba la Directiva N° 020-MINSA-DGSP-V.01: Sistema de Atención para el Tratamiento Antirretroviral en los Niños Infectados por el Virus de Inmunodeficiencia Humana. Lima, Peru, 2003.

16. Ministerio de Salud. Resolución Ministerial N° 567-2013/MINSA, que aprueba la NTS N° 102-MINSA/DGSP. V01: Norma Técnica de Salud para la Atención Integral y Tratamiento Antirretroviral de los Niños, Niñas y Adolescentes Infectados por el Virus de la Inmu.

17. Panel on Antiretroviral Therapy and Medical Management of HIV-Infected Children. Guidelines for the Use of Antiretroviral Agents in Pediatric HIV Infection. Available at http://aidsinfo.nih.gov/contentfiles/lvguidelines/pediatricguidelines.pdf. Accessed September 17, 2018.

18. Durrleman S, Simon R. Flexible regression models with cubic splines. Stat Med. 1989;8(5):551–61.

19. Li R, Hertzmark E, Spiegelman D. The SAS GLMCURV9 Macro. Boston, MA: Channing Laboratory; 2008.

20. Pan W. Akaike’s information criterion in generalized estimating equations. Biometrics. 2001;57(1):120–5.

21. Bangsberg DR. Less than 95% adherence to nonnucleoside reverse-transcriptase inhibitor therapy can lead to viral suppression. Clin Infect Dis. 2006;43(7):939–41.

22. Paterson DL, Swindells S, Mohr J, Brester M, Vergis EN, Squier C, et al. Adherence to protease inhibitor therapy and outcomes in patients with HIV infection. Ann Intern Med. 2000;133(1):21–30.

23. Valle E, Rodriguez C, Galea JT, Wong M, Kolevic L, Munoz M, et al. Understanding health-related behaviors among adolescents living with HIV in urban Peru; under review.

24. Galea JT, Wong M, Muñoz M, Valle E, Leon SR, Díaz Perez D, et al. Barriers and facilitators to antiretroviral therapy adherence among Peruvian adolescents living with HIV: A qualitative study. PLoS One. 2018;13(2):e0192791.

25. Quintana Y, Gonzalez Martorell EA, Fahy D, Safran C. A Systematic Review on Promoting Adherence to Antiretroviral Therapy in HIV-infected Patients Using Mobile Phone Technology. Appl Clin Inform. 2018;9(2):450–66.

26. Garrison LE, Haberer JE. Technological methods to measure adherence to antiretroviral therapy and preexposure prophylaxis. Curr Opin HIV AIDS. 2017;12(5):467–74.

27. MacPherson P, Munthali C, Ferguson J, Armstrong A, Kranzer K, Ferrand RA, et al. Service delivery interventions to improve adolescents’ linkage, retention and adherence to antiretroviral therapy and HIV care. Trop Med Int Health. 2015;20(8):1015–32.

28. Ridgeway K, Dulli LS, Murray KR, Silverstein H, Dal Santo L, Olsen P, et al. Interventions to improve antiretroviral therapy adherence among adolescents in low-and middleincome countries: A systematic review of the literature. PLoS One. 2018;13(1):e0189770.

29. IPSOS. Hábitos, usos y actitudes hacia el internet 2017. Accessible at: https://www.ipsos.com/es-pe/habitos-usos-y-actitudes-hacia-el-internet-2017. Accessed September 17, 2018.

30. Taggart T, Grewe ME, Conserve DF, Gliwa C, Roman Isler M. Social Media and HIV: A Systematic Review of Uses of Social Media in HIV Communication. J Med Internet Res 2015;17(11):e248.

31. Cao B, Gupta S, Wang J, Hightow-Weidman LB, Muessig KE, Tang W, et al. Social Media Interventions to Promote HIV Testing, Linkage, Adherence, and Retention: Systematic Review and Meta-Analysis. J Med Internet Res. 2017;19(11):e394.

32. Calderon Y, Cowan E, Nickerson J, Mathew S, Fettig J, Rosenberg M, et al. Educational Effectiveness of an HIV Pretest Video for Adolescents: A Randomized Controlled Trial. Pediatrics. 2011;127(5):911–6.

33. Bailey H, Cruz MLS, Songtaweesin WN, Puthanakit T. Adolescents with HIV and transition to adult care in the Caribbean, Central America and South America, Eastern Europe and Asia and Pacific regions. J Int AIDS Soc. 2017;20(0):21475.

34. Armstrong A, Nagata JM, Vicari M, Irvine C, Cluver L, Sohn AH, et al. A Global Research Agenda for Adolescents Living With HIV. J Acquir Immune Defic Syndr. 2018;78 Suppl 1:S16–21.

35. CIPHER. A global research agenda for adolescents living with HIV. 2017. Accessible at: https://www.iasociety.org/Web/WebContent/File/CIPHER_policy_brief_ado_EN.pdf. Accessed September 17, 2018.

36. Kanters S, Park JJH, Chan K, Socias ME, Ford N, Forrest JI, et al. Interventions to improve adherence to antiretroviral therapy: a systematic review and network metaanalysis. Lancet HIV. 2017;4(1):e31–40.

37. Govindasamy D, Meghij J, Negussi EK, Baggaley RC, Ford N, Kranzer K. Interventions to improve or facilitate linkage to or retention in pre-ART (HIV) care and initiation of ART in low-and middle-income settings – a systematic review. J Int AIDS Soc. 2014;17(1):19032.

38. Kim S-H, Gerver SM, Fidler S, Ward H. Adherence to antiretroviral therapy in adolescents living with HIV: systematic review and meta-analysis. AIDS. 2014;28(13).

39. Ministerio de Salud del Perú. Informe nacional sobre los progresos realizados en el país. Lima, Peru: 2014. Accessible at: http://files.unaids.org/en/dataanalysis/knowyourresponse/countryprogressreports/2014countries/PER_narrative_report_2014.pdf. Accessed September 17, 2018

40. Baker AN, Bayer AM, Viani RM, Kolevic L, Sim M-S, Deville JG. Morbidity and Mortality of a Cohort of Peruvian HIV-Infected Children 2003-2012. Pediatr Infect Dis J. 2018;37(6):564–9.

